# Monitoring the quality of eggs depending on the hens breeding systems by Raman spectroscopy

**DOI:** 10.1101/2022.02.16.480636

**Authors:** M. Kopec, H. Abramczyk

**Affiliations:** Lodz University of Technology, Institute of Applied Radiation Chemistry, Laboratory of Laser Molecular Spectroscopy, Wroblewskiego 15, 93-590 Lodz, Poland

**Keywords:** Raman spectroscopy, eggs, Raman biomarkers, chemometric methods

## Abstract

The aim of this study is show that Raman spectroscopy is a sensitive tool, which can be used for analyses eggs obtained in various hens housing system. This study provided a new and simple method for hens’ eggs breeding systems testing. Raman methods have potential to be applied by commercial inspections in food industry for verification indications placed on eggs. The development of functional technological methods to study the quality of eggs could be an interesting way to gain profitability for the food industry, in addition to improving the general conditions of public health. A label-free Raman method for detecting spectral changes in eggs from cage systems, barn systems, free range systems and ecological systems is presented. The most important advantage of Raman spectroscopy is the ability to identify biomarkers that help estimate the quality of eggs from various hens housing systems based on spectra typical for lipids, proteins and carotenoids. We have proved that ratios 1656/1004, 1656/1444, 1444/1520 and 1656/1156 can be used as universal biomarkers to distinguish eggs from various hens housing systems. Chemometric methods have shown that eggs from ecological systems and barn systems can be distinguished based on their vibrational properties.

**Graphical Abstract:** 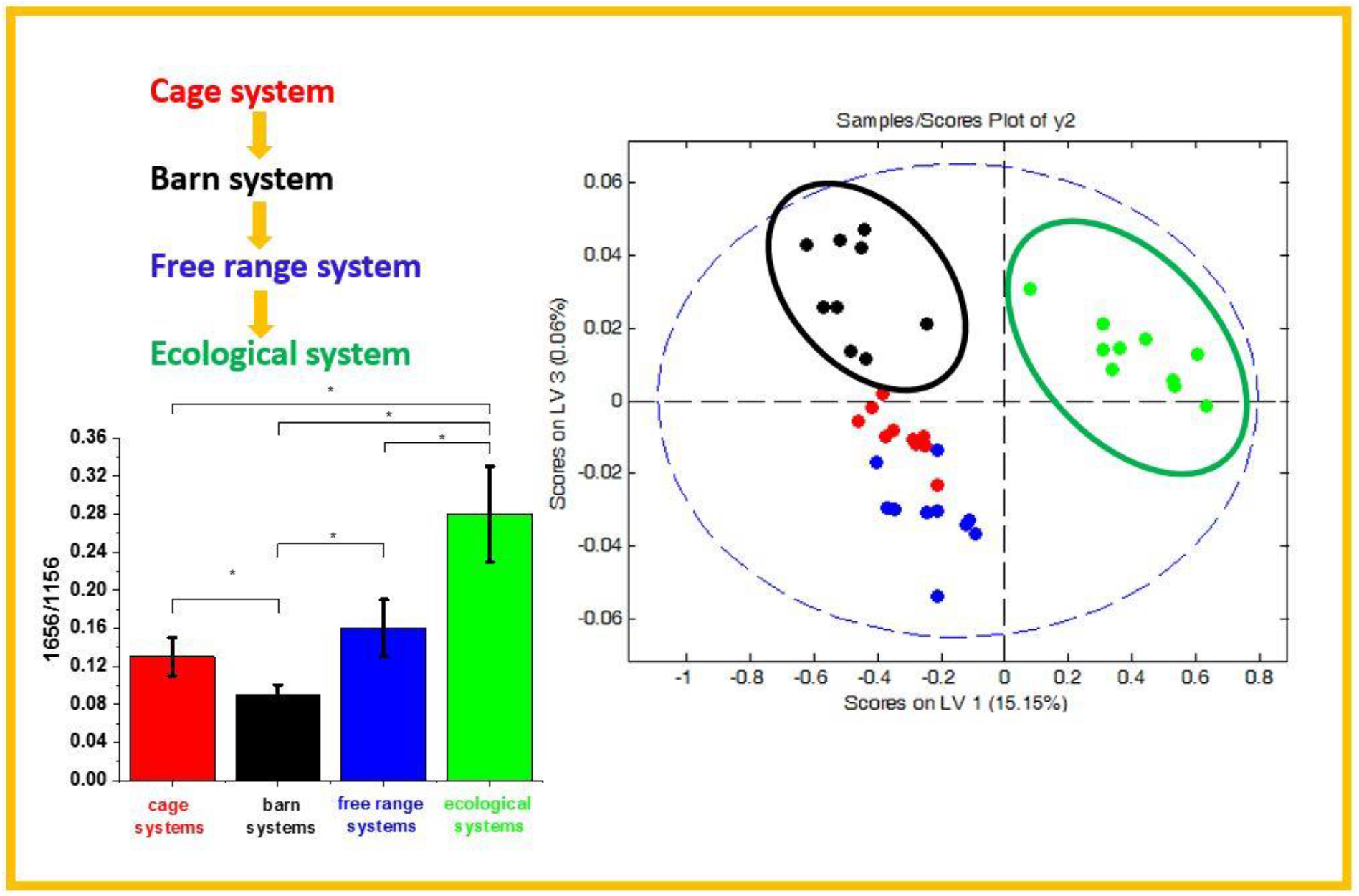

## Introduction

The human organism cannot produce antioxidants, so they must be incorporated by nutrition. Fruit and vegetables are rich source of antioxidants. But antioxidants are not only contained in plants. Animal products especially meat and eggs are sources of a variety of antioxidants. The vitamins A, C, E and D, carotenoids, lycopene, lutein and zeaxanthin are typical antioxidant substances in the human organism. Eggs especially the yolk contains a high concentration of different antioxidants such as carotenoids. [1]

Nowadays eggs are widely consumed worldwide. The egg industry is very important segment of the world food industry. Eggs are one of the most important components of human diet. [2] Nowadays, foods are not intended to only satisfy hunger but also to provide necessary nutrients for humans, to prevent nutrition-related diseases and improve physical and mental wellbeing of consumers. Eggs offer a moderate calorie source (about 150 kcal/100 g), a protein of excellent quality, great culinary versatility and low economic cost, which make eggs within reach to most of the population. Moreover eggs can be consumed throughout the word because there are no restrictions on religious grounds. [3] It is predicted that eggs consumption will be increasing in the future, because a lot of people change its consumption mode and food habits. [4]

Additionally the role of eggs in the therapy and prevention of chronic and infectious diseases have been reported. [5] [6] [7] [8] The role of antioxidants included in eggs especially carotenoids and zeaxanthin have been described in literature. The role of antioxidant in the prevention of macular degeneration, a leading cause of blindness in the elderly, and have been associated with lower risk of cataract extraction. [9] Epidemiology studies, such as that undertaken by D. Alexander et al. examine the effect of egg consumption on the risk of heart disease and stroke. They concluded that intake of up 1 egg daily may be associated with reduced risk of total stroke. [10] The role of egg yolk in antimicrobial activity has been studied by Kovas-Nolan et al. The researches have shown that immunoglobin Y-the component of egg yolk-inhibit infection and disease *in vitro* and *in vivo* symptoms of gastrointestinal pathogens. [7] Eggs also have anti-cancer activity. G. Sava et al. described the role of lysozyme in lung carcinoma. It has been reported that lysozyme is an anticancer agent which inhibit tumor formation and growth. [11]

Schematic structure of egg is presented in Figure 1. One can see from Figure 1 that typical egg consist of: cuticula, eggshell, chalaza, thin albumen, thick albumen, air cell, yolk and chalaza. [12]

**Fig. 1.**
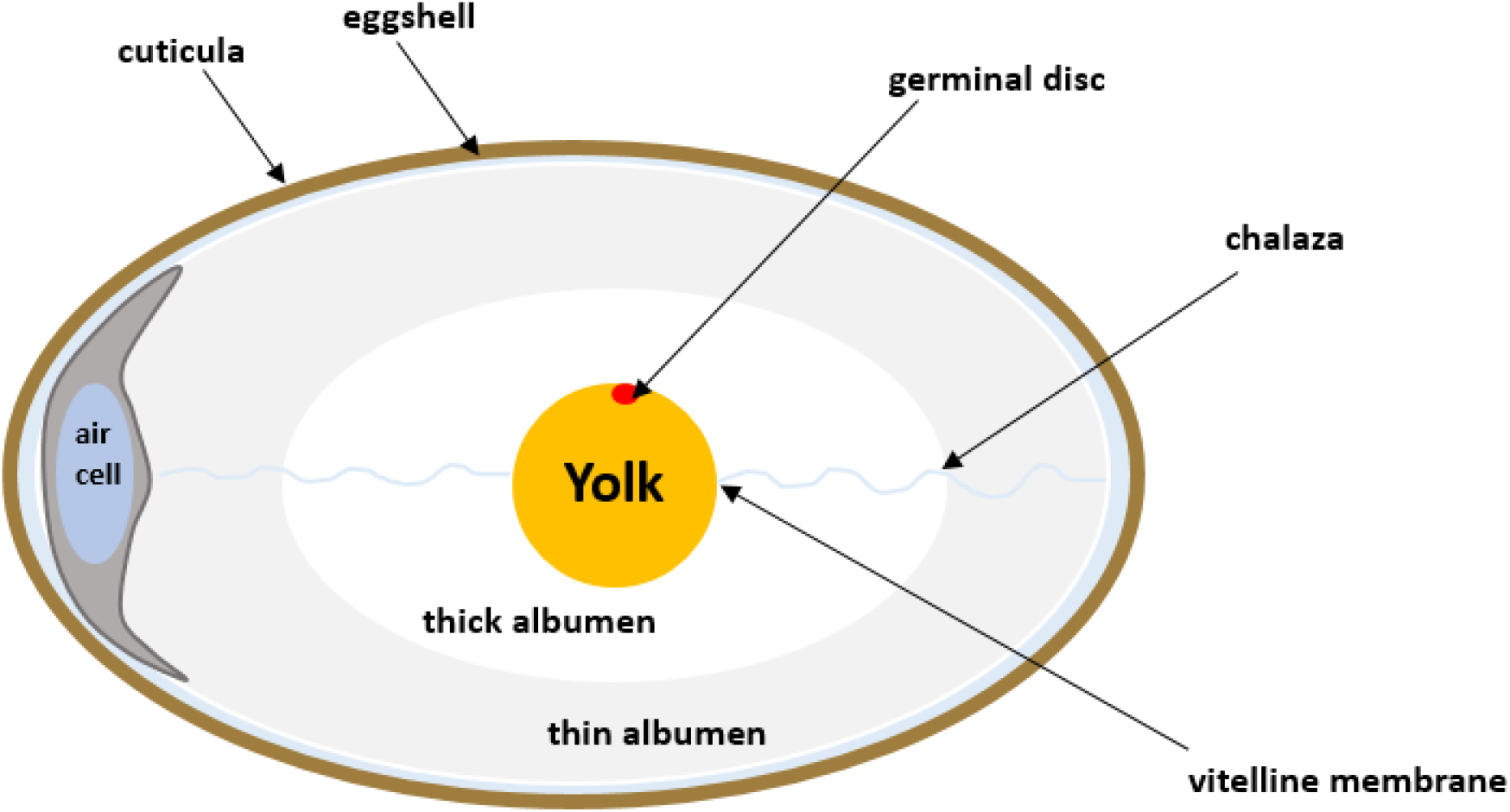
Schematic structures of egg

In general, an egg is covered by eggshell with a cuticle layer. Egg white is the major constituent of egg and occupies 60% of the whole egg. The yolk is an emulsified fluid and is central fraction of egg. The major components of yolk are proteins and lipids. The egg yolk lipids include triglycerides, phospholipids, cholesterol, cerebrosides or glycolipids, and some other minor lipids. [13] Egg yolk contains also pigments (carotenes and riboflavin) which are responsible for the colour of the yolk. Finally, egg yolk contains minerals in the greatest amount of phosphorus. [12]

Hen’s eggs are also source of proteins, so that their quality is critically important. Egg quality is determined based on many traits important for global egg production, and depends on many factors, such us: the diet, age of hens and the breeding systems. The breeding systems of hens have an important influence on the growth of hens, the prevention of diseases, and the yield and quality of eggs. There are several housing systems used in eggs production such as: cage systems, barn systems, free range systems, ecological systems. These systems have another environmental parameters of hens mainly temperature and humidity. [14] [15]

The quality of egg is depending on hens housing system. There are a lot of publication confirmed this observation. [16] [17] [18] [19] Van der Brand et al. investigated the quality of egg depending on hens housed conditions. They have noticed that yolk color was darker in the free-range eggs. [20] Other groups studied the heaviness of the eggs. They observed that eggs from free-range housing system are heavier and have higher albumen quality compared to cages system. [21]

In this manuscript we will focus on biochemical composition of hen’s egg from various hens housing conditions: cage systems, barn systems, free range systems and ecological systems. We will show that Raman spectroscopy can be used for bioanalytical characterization of hen’s eggs. Raman spectroscopy analysis of carotenoids, lipids and proteins profiles of eggs can be useful in determination of spectroscopic biomarkers for control the quality of eggs. The use of functional technological methods such as Raman spectroscopy to study the quality of eggs could be an interesting way to gain profitability for the food industry, in addition to improving the general conditions of public health.

## Results and discussion

To understand in more details the biochemical changes that occur in eggs we measured and analyzed the Raman spectra in various types of hens housing system. Raman spectroscopy was investigated to determine the biochemical composition of hen’s eggs in various breeding systems. Figure 1 presents the typical Raman spectra from hen’s yolk depending on hens housing conditions: cage systems (red line), barn systems (black line), free range systems (blue line) and ecological systems (green line) in fingerprint region (panel A, C and D) and in high frequency region (panel B). In Figure 1 one can see that Raman peaks typical for yolk eggs can be observed at: 955, 1004, 1156, 1189, 1265, 1299, 1444, 1520 and 1656 cm^-1^ in fingerprint region and at: 2666, 2849, 2898, 2910, 2921 and 3002 cm^-1^ in high frequency region. The Raman spectrum contains biochemical information about the sample. The detailed assignments of Raman peaks observed in Fig. 1 are presented in Table 1.

**Table 1.** The vibrational assignment of the Raman spectra [22]

| Wavenumber [cm<sup>-1</sup>] | Tentative assignments |
| --- | --- |
| 955 | CH=CH bending |
| 1004 | proteins |
| 1156 | carotenoids |
| 1189 | C-C<sub>6</sub>H<sub>5</sub> Phe, Trp |
| 1265 | Fatty acids, =C-H bend |
| 1299 | CH<sub>2</sub> deformation of lipids |
| 1444 | Lipids, fatty acid |
| 1520 | carotenoids |
| 1656 | unsaturated fatty acids, triglycerides (C=C) str, |
| 2666 | an overtone of carotenoids |
| 2849 | Fatty acids, triglycerides, C-H<sub>2</sub> symmetric stretching |
| 2898 | CH<sub>2</sub> antysymmetric stretch of lipids and proteins |
| 2910 | proteins |
| 2921 | proteins |
| 3002 | lipids |

Comparing the spectra from Fig 1 more detailed biochemical information about eggs from various hen living system can be obtained. One can see that main biochemical differences are observed in fingerprint region. For better monitoring changes in yolk from hens living in various housing conditions we decided to enlarge the ranges 1400-1600 cm^-1^ and 1600-1750 cm^-1^ (Fig. 1 C, D). A detailed examination of Figure 1 panels (C, D) shows some differences. The most relevant differences presented on Fig 1 are related to the vibrations of fatty acid and lipids (1444 cm^-1^), unsaturated fatty acids, triglycerides (1656 cm^-1^) and (an overtone of carotenoids) 2666 cm^-1^.

Firstly, the intensity of peaks 1444 cm^-1^ corresponding to fatty acids and lipids is much stronger in ecological systems (green line) than in barn systems (black line). The amount of fatty acids and lipids are different depending on hens breeding systems. This observation confirms that eggs from ecological systems in contrary to the barn systems have higher concentration of lipids and fatty acid. Secondly one can see differences at 1656 cm^-1^ corresponding to unsaturated fatty acids, triglycerides (C=C) stretching. The band at 1656 cm^-1^ has the highest intensity for ecological systems and the lowest intensity for eggs from barn systems. Spectacular difference is also observed at 2666 cm^-1^. This band attributed to carotenoids is much stronger in eggs from barn system (black line) than in eggs from ecological system (green line).

Based on the results shown in Table 1 and frequencies highlighted in Fig. 1 we have chosen the following Raman bands to compare: 1004, 1156, 1444, 1520 and 1656 cm^-1^. To identify the changes occurring in Raman spectra for yolk eggs in various hens housing conditions we decided to calculate ratios: 1656/1004, 1656/1444, 1444/1520 and 1656/1156. Table 1 and Figure 2 shows the comparison of Raman band intensity ratios for four different hens housing system: cage system, barn system, free range system and ecological system.

**Fig 2.**
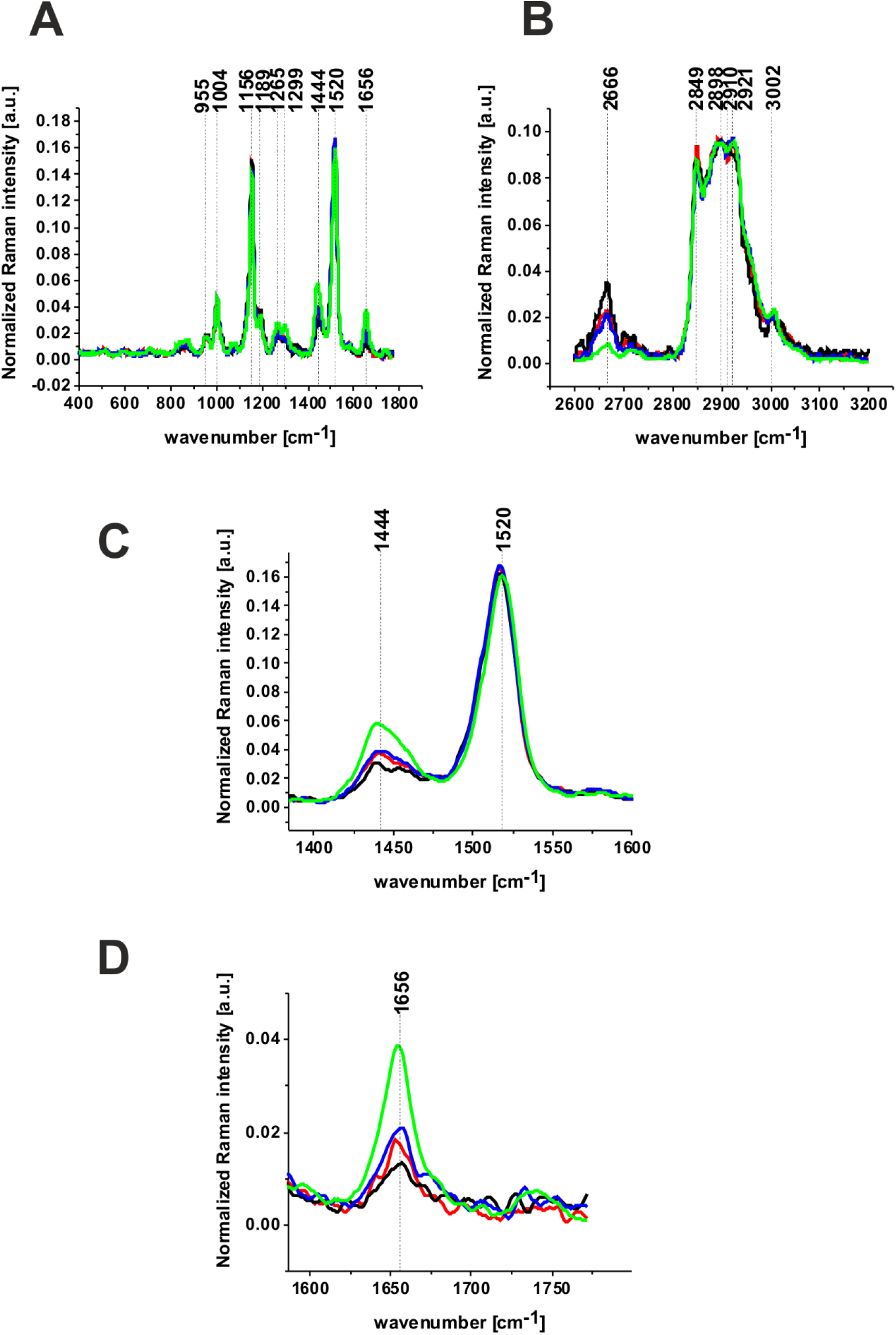
Raman spectra of yolk hen eggs from cage systems (red line), barn systems (black line), free range systems (blue line) and ecological systems (green line)

To better understand differences, in yolk eggs from hens living in various conditions, obtained from Raman spectroscopy we will concentrate on the statistical analysis. We analyses the data using Origin. To check the statistical difference between the populations we used the one-way Anova Tukey test. p-Values ≤ 0.05 were accepted as statistically significant. Results which are statistically significant we marked with an asterisk.

One can see from Fig. 3 and Table 2 that presented above ratios are useful to discriminate the type of hens living system. Analyzing the ratios for vibrations of fatty acid, lipids, unsaturated fatty acids and triglycerides non-negligible differences are visible for the eggs from ecological system in comparison to the eggs from cage system, barn system and free range system. It is evident from Figure 3 that all ratios are the highest for yolk eggs from hens live in ecological systems. Additionally, the ratios 1656/1004, 1444/1520, 1656/1156 and 1656/1444 are the smallest for eggs from hens live in barn systems. In the view of the results presented in Figures 2, 3 and table 2 it is evident that the Raman biomarkers 1656/1004, 1444/1520, 1656/1156 and 1656/1444 measuring contribution of unsaturated fatty acids and triglycerides/proteins; lipids and fatty acids/carotenoids; unsaturated fatty acids and triglycerides/carotenoids, unsaturated fatty acids and triglycerides/lipids and fatty acids in the eggs correlates with hen housing systems.

**Table 2.** Raman intensity ratios 1656/1004, 1656/1444, 1444/1520, 1656/1156 for yolk eggs from hens live in cage systems, barn systems, free range systems and ecological systems.

| Ratio | Cage Systems |  | Barn Systems |  | Free Range Systems |  | Ecological Systems |  |
| --- | --- | --- | --- | --- | --- | --- | --- | --- |
|  | Mean | SD | Mean | SD | Mean | SD | Mean | SD |
| 1656/1004 | 0,4 | 0,07 | 0,32 | 0,07 | 0,47 | 0,07 | 0,8 | 0,1 |
| 1656/1444 | 0,5 | 0,04 | 0,46 | 0,05 | 0,55 | 0,05 | 0,63 | 0,13 |
| 1444/1520 | 0,23 | 0,02 | 0,19 | 0,02 | 0,24 | 0,03 | 0,36 | 0,05 |
| 1656/1156 | 0,13 | 0,02 | 0,09 | 0,01 | 0,16 | 0,03 | 0,28 | 0,05 |

**Fig 3.**
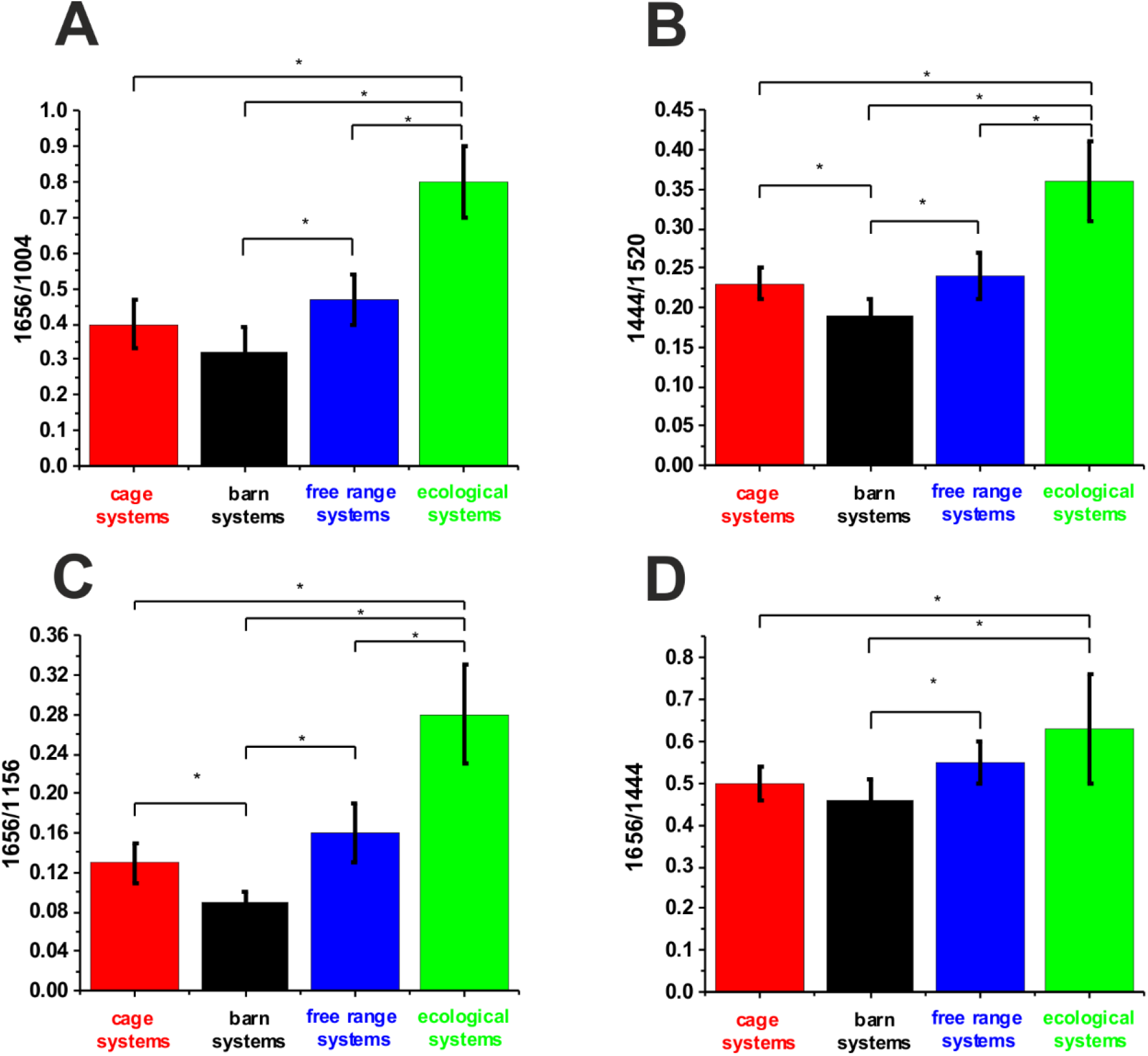
The Raman intensity ratios and SD for 1656/1004 (A), 1444/1520 (B), 1656/1156 (C), 1656/1444 (D). The statistically significant results have been marked with an asterix.

Additionally, to graphical visualize the differences between hens living systems in various conditions we performed statistical analysis using chemometric method PLS-DA. Figure 4 shows the PLS-DA score plot corresponding to the Raman spectra of yolk eggs from hens living in various conditions.

**Fig 4.**
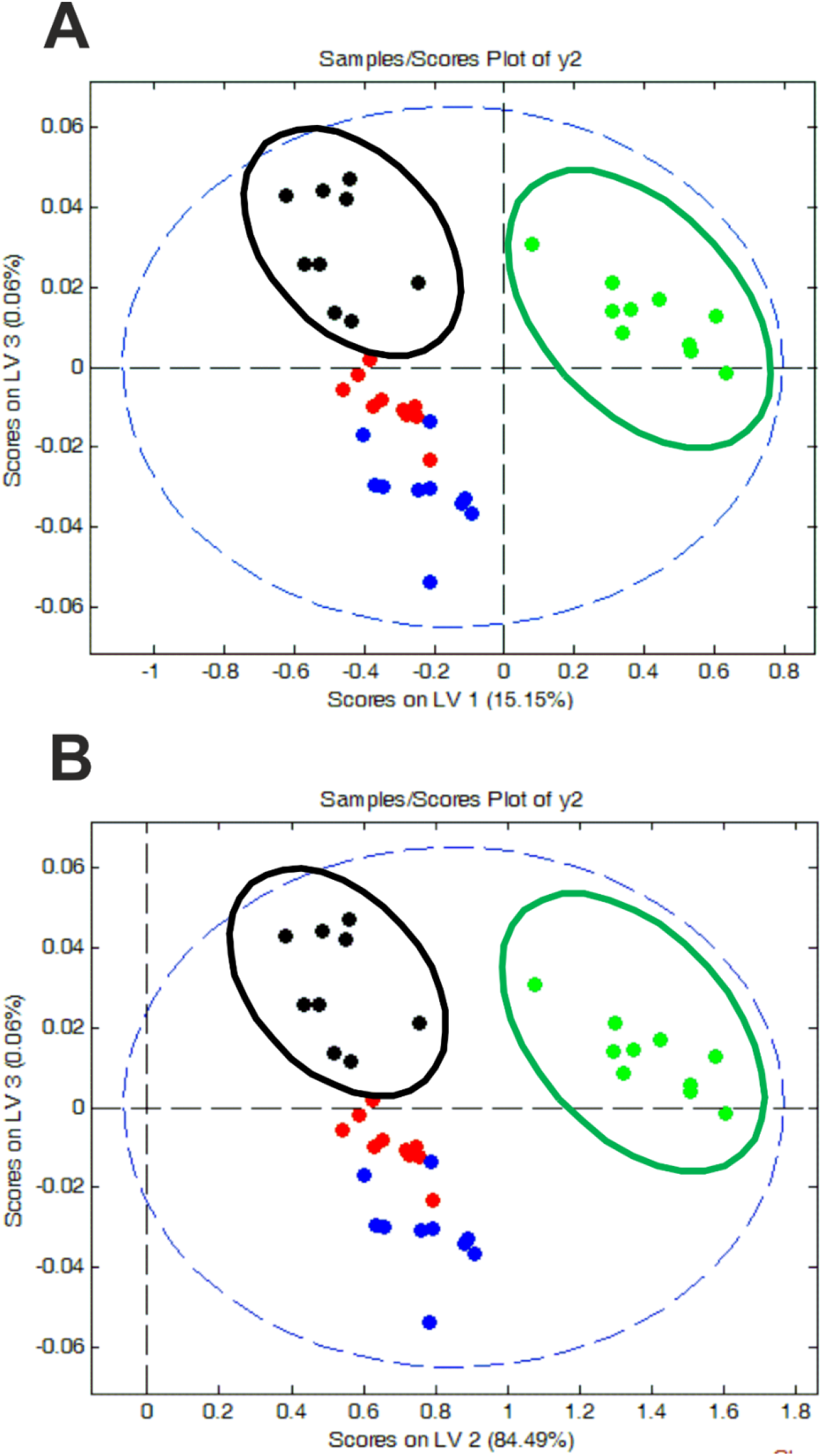
PLSDA score plot for the Raman spectra for: cage systems (red circle), barn systems (black circle), free range systems (blue circle) and ecological systems (green circle)

By plotting the component scores, similarities between the samples can be revealed. The plots presented on Figure 4 evidently confirm the differences between yolk hen’s eggs from various housing conditions. The differences and similarities are visible by grouping the results into separate clusters. Raman spectra for ecological system (green circle) are the most separated from the rest. Moreover, one can see that the Raman spectra for barn system (black circle) are also separated. The spectra for ecological system and barn system are grouped in the upper area while the spectra from free range systems (blue circle) and cage systems (red circle) are grouped in the lower area of the plot. One can see that Raman spectra for cage systems (red circle) and for free range systems (blue circle) are mixed. This result was presented also in Figure 1.

To understand the molecular information contained in the LV1, LV2 and LV3 we used the loading plots presented in Fig. 5 that reveal the most important characteristic features in the Raman spectra.

**Fig 5.**
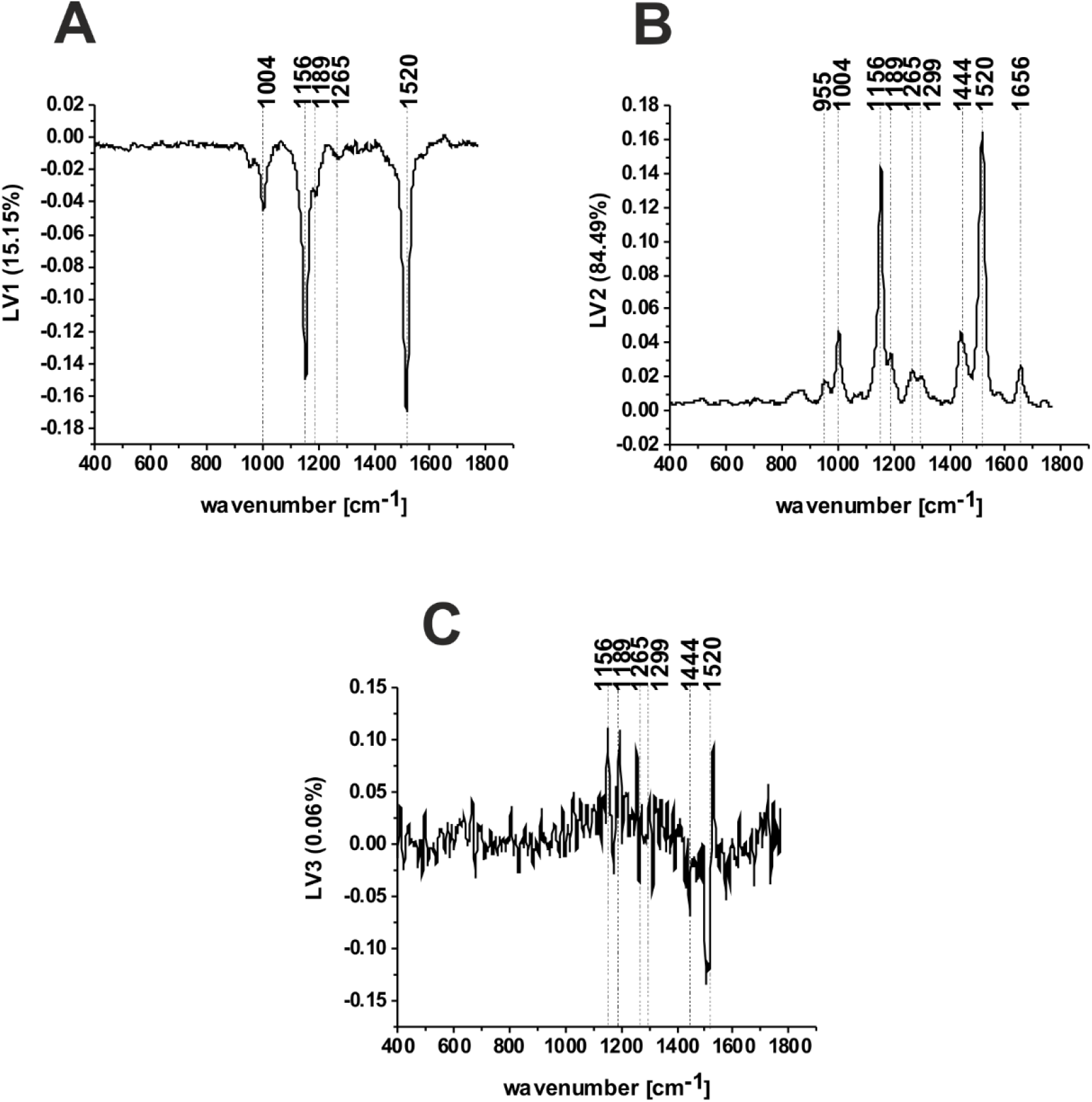
Loading plots of LV1 (A), LV2 (B), LV3 (C) versus wavenumber [cm^-1^] obtained from PLS-DA analysis.

Figure 5 shows the loading plots of LV1, LV2 and LV3 as a function of the wavenumber. Once can see that the loading plots show the most pronounced changes around the characteristic Raman peaks of carotenoids and proteins.

Once can see from Fig. 5 that the first LV has the contribution of 15.15%, LV2 the contribution of 84.49% and LV3 the contribution of 0.06% to variance, respectively. LV1 and LV2 give the dominant account for the maximum variance in the data. The first latent variable LV1 is presented in Fig. 5 A. The most characteristic minima in the loading plot are at 1004 cm^-1^ (proteins), 1156 cm^-1^ (carotenoids) and at 1520 cm^-1^ (carotenoids). The second latent variable LV2 (Fig. 5B), gives the highest contribution to variance, reaches its maxima at 1004 (proteins), 1156 (carotenoids), 1444 (Lipids, fatty acid), and 1656 cm^-1^ (unsaturated fatty acids, triglycerides (C=C) stretching). The third latent variable presented in Fig. 5C has only % to variance.

To access the potential of Raman spectroscopy for control the quality of eggs we calculated sensitivity and specificity. Table 3 presents the value of sensitivity and specificity obtained from PLS-DA method. The high values for sensitivity and specificity highlight the importance of Raman spectroscopy as a new tool for eggs quality control.

**Table 3.** The value of sensitivity and specificity for calibration and cross validation procedure from PLS-DA analysis

|  | cage<br>systems | barn<br>systems | free range<br>systems | ecological<br>systems |
| --- | --- | --- | --- | --- |
| Sensitivity (calibration) | 1.0 | 1.0 | 1.0 | 1.0 |
| Specificity (calibration) | 0.586 | 0.967 | 0.966 | 1.0 |
| Sensitivity (cross validation) | 1.0 | 0.889 | 0.8 | 1.0 |
| Specificity (cross validation) | 0.517 | 0.933 | 0.897 | 1.0 |

One can see from Table 3 that calculated sensitivity and specificity are the highest for eggs from ecological system. The lowest value of specificity in calibration and cross validation is obtained for cage system and takes values 0.586 and 0.517, respectively. The high values for sensitivity and specificity highlight the importance of Raman spectroscopy as a new method in the egg quality control process.

## Conclusions

In this study we have shown that using Raman spectroscopy there is possibility to determine the eggs from various hens housing system. It was proven that Raman spectroscopy is effective tools for monitoring and analyzing the vibrations of functional groups in eggs. The results presented in this paper demonstrate that using Raman spectroscopy we can control the amount of carotenoids, lipids and proteins in eggs from various hens housing system. We showed that the ratios 1656/1004, 1656/1156 and 1444/1520 play a crucial role as a Raman biomarker of the verification hens housing system. The analysis of the intensity of the lipids/proteins/carotenoids bands shows that these classes of compounds can classify the analyzed samples into hens housing systems. PLS-DA method confirm that for the Raman spectra the sensitivity for every hens housing system equal 100%. The specificity equal 59%, 97%, 97% and 100% for cage systems, barn systems, free range systems and ecological systems, respectively. The results presented in this work demonstrate that Raman spectroscopy is powerful method that will bring an important contribution to the verify the designation on eggs relating to the hens housing system and control the quality of eggs.

## Materials and methods

### 1. Eggs

The fresh eggs were obtained from hens in various housing conditions such as: cage systems, barn systems, free range systems and ecological systems. To prepare eggs for the Raman spectroscopy measurements the eggshells were opened, and the yolk was separated from the rest of the ingredients. Afterwards, the drop of yolk was put on CaF_2_ windows. For every hen housing conditions, we analyzed 10 eggs.

### 2. Raman spectroscopy

All Raman spectra presented in this manuscript were recorded using a confocal Raman microscope (WITec (alpha 300 RSA+) using a 50 μm core diameter fiber, 532 nm excitation line, an imaging spectrograph/monochromator (Acton-*S*P-2300i), a CCD camera (Andor Newton DU970-UVB-353) and an Ultra High Throughput Spectrometer. During Raman measurement we have used 40x dry objective and the average laser excitation 10mW. Every day before Raman measurements the confocal system was calibrated using silicon plate (520.7 cm^−1^). The Raman spectra for each egg were recorded with integration time 1 sec and 10 accumulations in the fingerprint region and in the high frequency region. Each Raman spectrum was preprocessed using the WITec Control/Project Four 4.1 package. Briefly, cosmic rays’ removal, smoothing and background corrections. More detailed description of equipment is available in our previous papers. [23–25]

### 3. Statistical analysis

Data presented in this manuscript were normalized using Origin software (normalization model: divided by norm). All results regarding the analysis of the intensity of the Raman spectra as a function of hens housing type are presented as the mean ± SD, where p less than 0.05 (SD—standard deviation, p—probability value). A significance level of less than 0.05 was used for all statistical analyses (marked with an asterisk). The average spectra, ratios and standard deviations were also calculated in Origin. One-way Anova Tukey tests are also prepared in Origin.

PLS-DA analysis was performed using MATLAB (MathWorks, USA) with PLS-Toolbox (EigenvectorResearch Inc., USA). Detailed information of chemometric methods are available elsewhere. [23–25]

## Founding

This research was funded by Statutory activity 2022: 501/3-34-1-1

